# Dry surface biofilm formation by *Candida auris* facilitates persistence and tolerance to sodium hypochlorite

**DOI:** 10.1101/2023.10.02.560537

**Authors:** Alicia Ware, William Johnston, Christopher Delaney, Mark Butcher, Gordon Ramage, Lesley Price, John Butcher, Ryan Kean

## Abstract

*Candida auris* is an enigmatic fungal pathogen, recently elevated to the critical priority group of pathogens by the World Health Organization. Of key concern is its ability to cause outbreaks within intensive and chronic care units, facilitated through its environmental persistence. We investigated the susceptibility of phenotypically distinct *C. auris* isolates to sodium hypochlorite (NaOCl) disinfection, and the subsequent role of biofilms in surviving disinfection using a dry-surface biofilm (DSB) model and transcriptomic profiling. Planktonic cells were tested for susceptibility to NaOCl in suspension, with biofilm formation using the DSB model consisting of consecutive 48 hr cycles with/without media across a 12-day period, assessed using viable counts, biomass assays, and microscopy. Disinfection efficacy was assessed using clinically relevant protocols of 500-1000ppm for 1-5min. RNA-sequencing was performed on untreated DSBs in comparison to planktonic cells. Isolates were found to be sensitive to NaOCl planktonically at concentrations ≤62.5 ppm, and grew robust biofilms using the DSB protocol. Biofilms developed tolerance to all NaOCl treatment parameters, with only 2-4 log_10_-reductions in viable cells observed at highest concentrations. Transcriptomics identified ABC transporters and iron acquisition pathways as strongly upregulated in DSBs relative to planktonic cells. Our novel findings have optimised a DSB protocol in which *C. auris* biofilms can mediate tolerance to adverse conditions such as NaOCl disinfection, suggesting a lifestyle through which this problematic yeast can environmentally persist and transmit. Mechanistically it has been shown for the first time that upregulation of small-molecule and iron transport pathways are potential facilitators of environmental survival.

**IMPORTANCE:** *Candida auris* is a pathogenic yeast that has been responsible for outbreaks in healthcare facilities across the globe, predominantly affecting vulnerable patients. This organism displays a concerning ability to persist within the healthcare environment that is likely facilitated by attaching onto surfaces and developing protective microbial communities knows as biofilms. These communities allow cells to survive and tolerate disinfection with bleach. In this study, we show that *C. auris* forms robust biofilms on surfaces which promote survival up to 12 days, even with prolonged drying periods. We also demonstrate that development of these biofilms over time significantly reduces the efficacy of hypochlorite disinfection. By investigating the molecular mechanisms of biofilms, we have shown that these biofilms express efflux pumps, which may actively remove hypochlorite molecules from cells, allowing them to tolerate disinfection, and that uptake of iron from the external environment is also important for survival of these communities.

## INTRODUCTION

In little over a decade, *Candida auris* has emerged as a significant nosocomial threat, responsible for outbreaks across the globe, and most recently exacerbated by the Covid-19 pandemic (1). In late 2022 the World Health Organization highlighted *C. auris* as one of four fungal pathogens in the Critical Priority group, owing to its intrinsic resistance to certain antifungal agents and ability to cause life-threatening infections resulting in unacceptably high mortality rates (1). *C. auris* readily forms biofilms on biotic and abiotic surfaces *in vitro* (2, 3), a survival mechanism which is strongly suspected in persistence within the healthcare setting (4). These communities of cells have routinely been shown to tolerate increased concentrations of all three classes of antifungals, facilitated through drug sequestration by the extracellular matrix (ECM) and upregulation of efflux pumps *CDR1* and *MDR1* (5, 6). *In vitro* evidence also suggests that biofilms also facilitate skin colonisation, with *C. auris* displaying enhanced adhesive capacity on both human and porcine skin models in comparison to *C. albicans* (3). Although environmental contamination is a likely reservoir for transmission of *C. auris* between hosts (7), the role biofilms may play in this process is less well-studied.

One concept that links environmental contamination with hospital acquired infections is biofilm formation on dry surfaces (8). Most extensively studied in the bacterial pathogen *Staphylococcus aureus*, these biofilms have been identified *in situ* on fomites within high contact areas such as patient folders, keyboards and sanitising bottles (8). *In vitro* models have shown that these communities can withstand environmental stressors such as temperature (9), physical removal, and disinfection (10). Given the capacity for *C. auris* to survive on various substrates for extended periods of time under both dry and hydrated conditions (11), combined with reported clinical transmission facilitated by reusable equipment such as temperature probes (12), a function for biofilms on dry surfaces in nosocomial outbreaks is highly plausible.

Cleaning and disinfection of environmental contamination is therefore of key importance in preventing transmission and outbreaks of *C. auris.* While a number of studies have demonstrated concentration- and contact time-dependent efficacy of biocides with various active agents when tested against *C. auris* in suspension (13–16), biofilm testing has reported decreased efficacy of compounds such as sodium hypochlorite (NaOCl) (2). Chlorine-based, oxidising agents such as NaOCl are widely regarded as the gold standard for disinfection of surfaces and equipment, owing to a wide spectrum of activity (17). Given the knowledge gap in understanding survival strategies of *C. auris* in response to biocidal antimicrobial agents, we herein investigated the function and mechanism of biofilms mimicking those formed on nosocomial dry surfaces in response to sodium hypochlorite disinfection. Here we report for the first time that DSBs of *C. auris* clinical isolates from outbreak-associated clades develop tolerance to NaOCl over subsequent cycles, facilitated by upregulation of efflux pumps and iron scavenging mechanisms.

## RESULTS

### Sensitivity of planktonic cells to sodium hypochlorite

Sensitivity to NaOCl has previously been reported for planktonic cultures of C. auris clinical isolates using a variety of methods (11, 14, 15, 18–20). Intrinsic sensitivity to the chemical activity of NaOCl was confirmed in planktonic cells of six *C. auris* isolates of different phenotypes. The activity of NaOCl following 24 h co-culture was similar across the six isolates (**Fig. 2A**). The MIC_50_ was shown to be 31.25 ppm for all isolates except NCPF8990 and NCPF8991, both of which had MIC_50_ of 62.5 ppm (**Fig. 2B**). No growth was observed amongst isolates for concentrations of NaOCl above 62.5 ppm, which is well below the concentration recommended for routine cleaning (21).

**Fig. 1.**
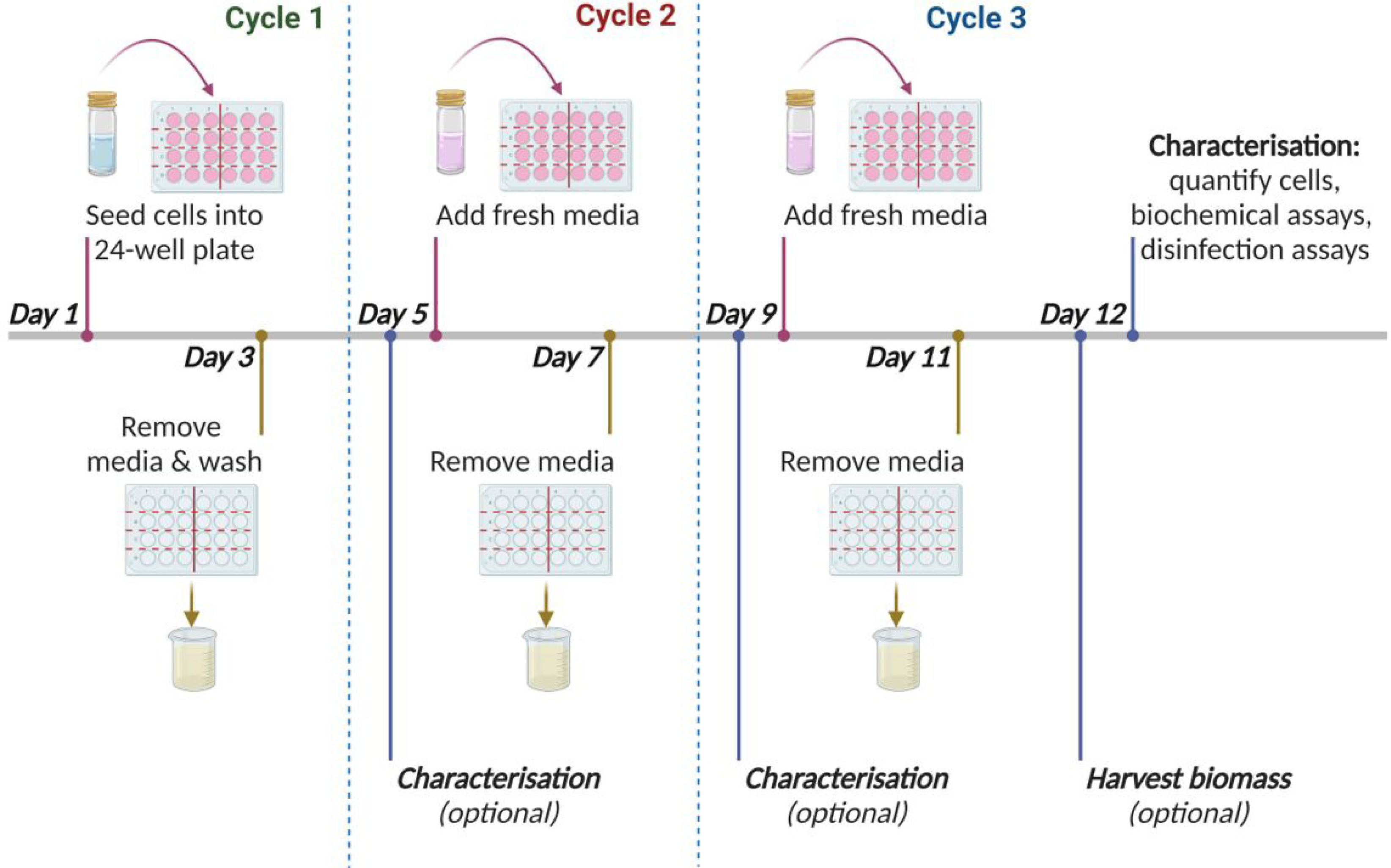
The dry surface biofilm protocol. Isolates of *C. auris* were grown as dry surface biofilms (DSB) in appropriate growth medium over three cycles, each consisting of 48 h ‘wet’ phase where media was present, followed immediately by 48 h ‘dry’ phase wherein all media was removed from wells. Plates were incubated at room temperature (22 – 24.5°C) at all times. Biofilms were assayed or treated at the conclusion of each dry phrase to determine the impact of each cycle on biofilm characteristics. Red lines indicate the beginning of each wet phase; brown lines indicate the beginning of each dry phase; blue dashed lines indicate the end of each cycle, at which time point characterisation assays were conducted (blue solid lines). Figure created using <u>Biorender.com.</u>

**Fig. 2.**
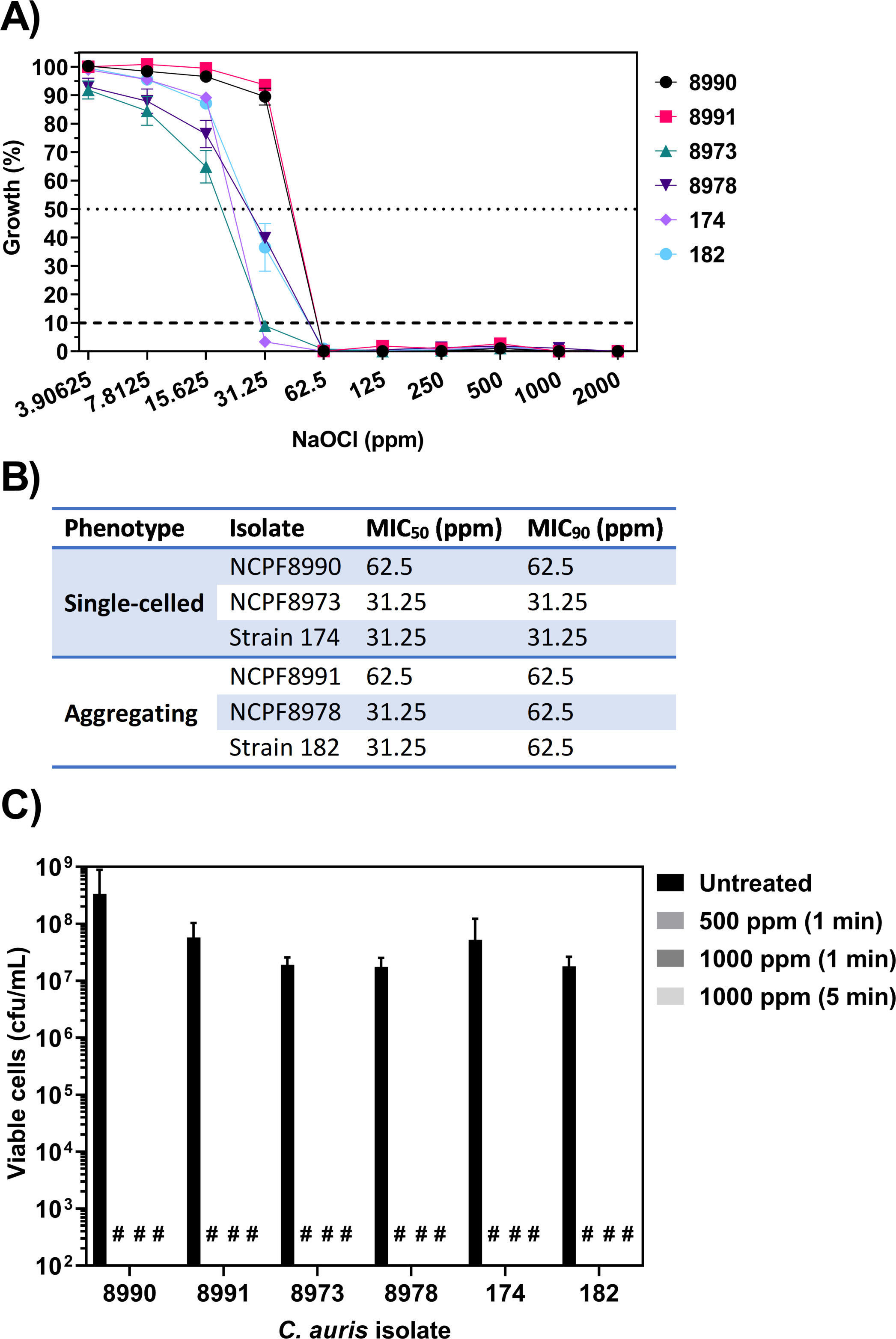
Susceptibility testing of *Candida auris* clinical strains against sodium hypochlorite. (A) Planktonic cells were co-cultured with increasing concentrations of sodium hypochlorite (NaOCl) for 24 h. Growth of exposed cells was measured using absorbance at 530 nm and normalised to untreated cells for each strain. Dashed lines indicate growth below 50% and 10% of the positive control. (B) The minimum concentration of NaOCl required to inhibit 50% (MIC_50_) and 90% (MIC_90_) of cellular growth relative to untreated cells, was determined for each isolate. (C) Planktonic cells were treated with sodium hypochlorite (NaOCl) at 500 – 1000 ppm available chlorine for contact time of 1 or 5 min, or sterile water (untreated) for 5 min, before neutralisation of active agents. Data represent the mean (± SEM) for n=3 experiments; # represents no cfu detected.

The PMIC method for determining susceptibility does not accurately reflect all circumstances under which NaOCl is used as a disinfectant. We therefore evaluated efficacy of NaOCl as a surface disinfectant when applied for 1 or 5 min, at manufacturer-recommended concentrations of 500-1000 ppm active chlorine, and with neutralisation Na_2_S_2_O_3_ to standardise contact times. Treatment with NaOCl resulted in a ≥6-log_10_ reduction in viable cells compared to the untreated cells across all six isolates, with no viable cells recovered after any treatment (**Fig. 2C**).

### Characterising dry surface biofilm formation by *C. auris* isolates

The DSB model has previously been used simulate growth of *C. auris* on stainless steel surfaces following contamination with biological material (22). We screened the six different isolates for ability to form biofilms on polystyrene surfaces under DSB conditions. Isolates were previously characterised as high-, moderate-, or low-biofilm forming isolates using a 24 h model (HBF, MBF, and LBF, respectively; **Fig. S1**). All isolates developed biofilms using the DSB protocol, with viable cells and varying degrees of biomass detected in all strains across all time points (**Fig. 3A,B**). The MBF isolates NCPF8973 and NCPF8978 demonstrated greatest number of viable cells across all timepoints which suggests they may be better suited to DSB formation than the other isolates. These two isolates were therefore selected for subsequent experiments. Using SEM, we observed dense, multi-layered biofilms of isolate NCPF8973 across most areas of the substrate following the first DSB cycle, with other areas characterised by less dense monolayers of cells (**Fig. 3Ci**). Higher magnification revealed presence of ECM on the surface of some cells, with limited connections to other cells. By the final cycle, the substrate was thickly covered with higher-density, multi-layered biofilms with interconnecting ECM covering all cells (**Fig. 3CiI**).

**Fig. 3.**
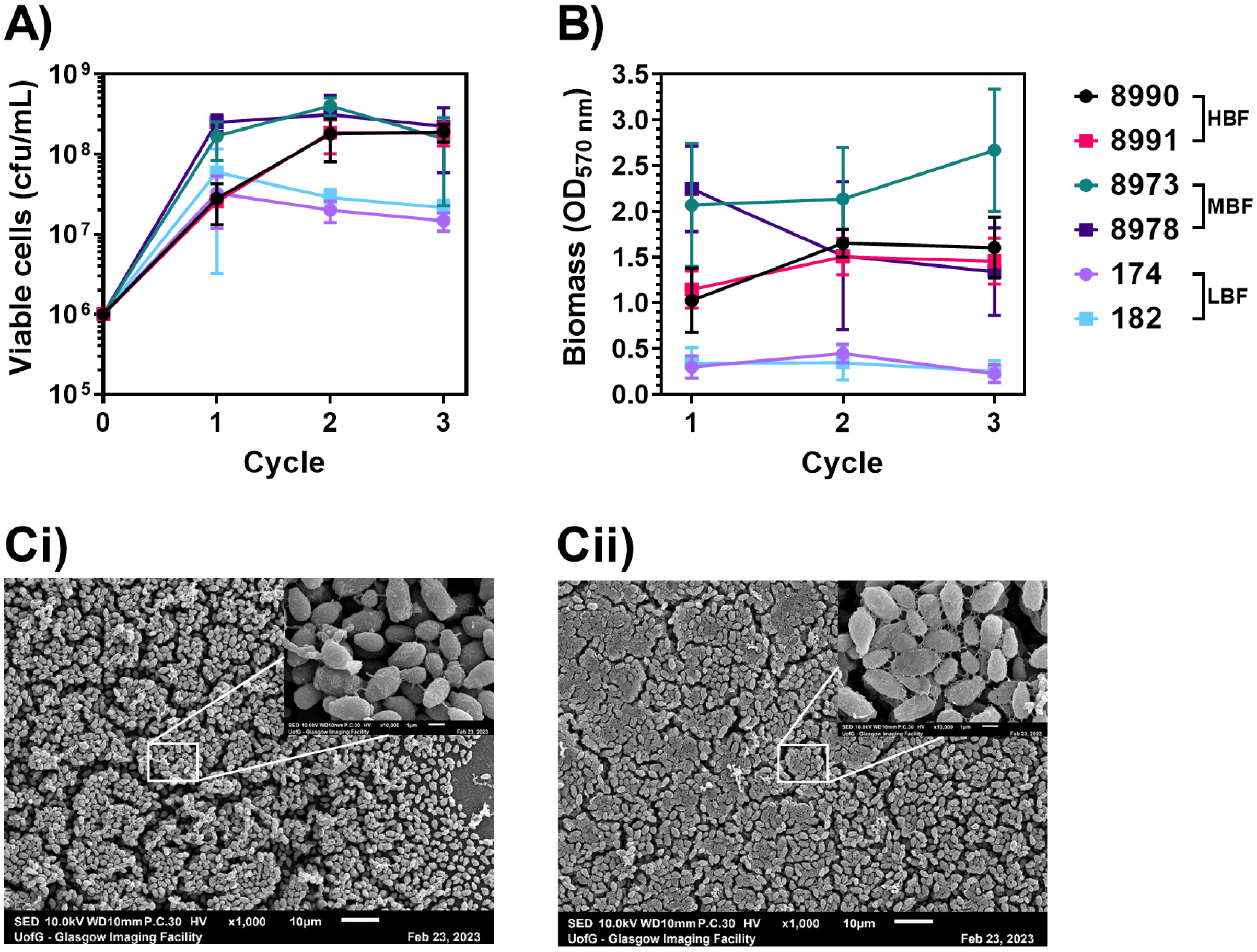
*Candida auris* form multilayer biofilms under dry surface conditions. (A, B) Clinical isolates of *C. auris* were standardised to 1×10^6^ cells/mL and grown as biofilms over one, two or three successive cycles of the DSB growth protocol. Viable cells (A) and biomass (B) were quantified at the conclusion of each cycle using plate counts and crystal violet assay, respectively. (C) Dry surface biofilms of single-celled isolate NCPF8973 were grown on Thermanox coverslips over one (Ci) or three (Cii) successive cycles of wet/dry conditions. Biofilm ultrastructure was imaged using scanning electron microscopy at x1000 and x10000 (inset).

### Development of NaOCl tolerance in dry surface biofilms

Given that *C. auris* biofilms display tolerance to antifungals and other small molecules (6), we further investigated the ability of *C. auris* DSBs to withstand chemical disinfection with NaOCl. Across the three cycles, disinfection with NaOCl resulted in statistically significant reductions in viable cells for both isolates, regardless of the contact time and concentration of hypochlorite (p<0.01; **Fig. 4**). Each subsequent cycle, however, resulted in improved survival of disinfectant-treated cells, evidenced by a decrease in log_10_-reductions in viable cells for each treatment. Disinfection efficacy of 1 min at 500 ppm was most affected by subsequent growth of NCPF8973 DSBs, as the mean log_10_-reduction in cfu/mL decreased from 4.5 to 1.2 from the first to the final cycle (**Fig. 4A**). In contrast, the most affected treatment for NCPF8978 was 1000 ppm for 5 min, with a decrease in mean reduction in log_10_-cfu/mL from 6.7 to 3.0 across the three cycles (**Fig. 4B**). Furthermore, for the first two cycles, disinfection with 1000 ppm for 5 min resulted in complete eradication of cells in some samples, whereas by the final cycle all biofilms contained viable cells following treatment (**Fig. 4A,B**).

**Fig. 4.**
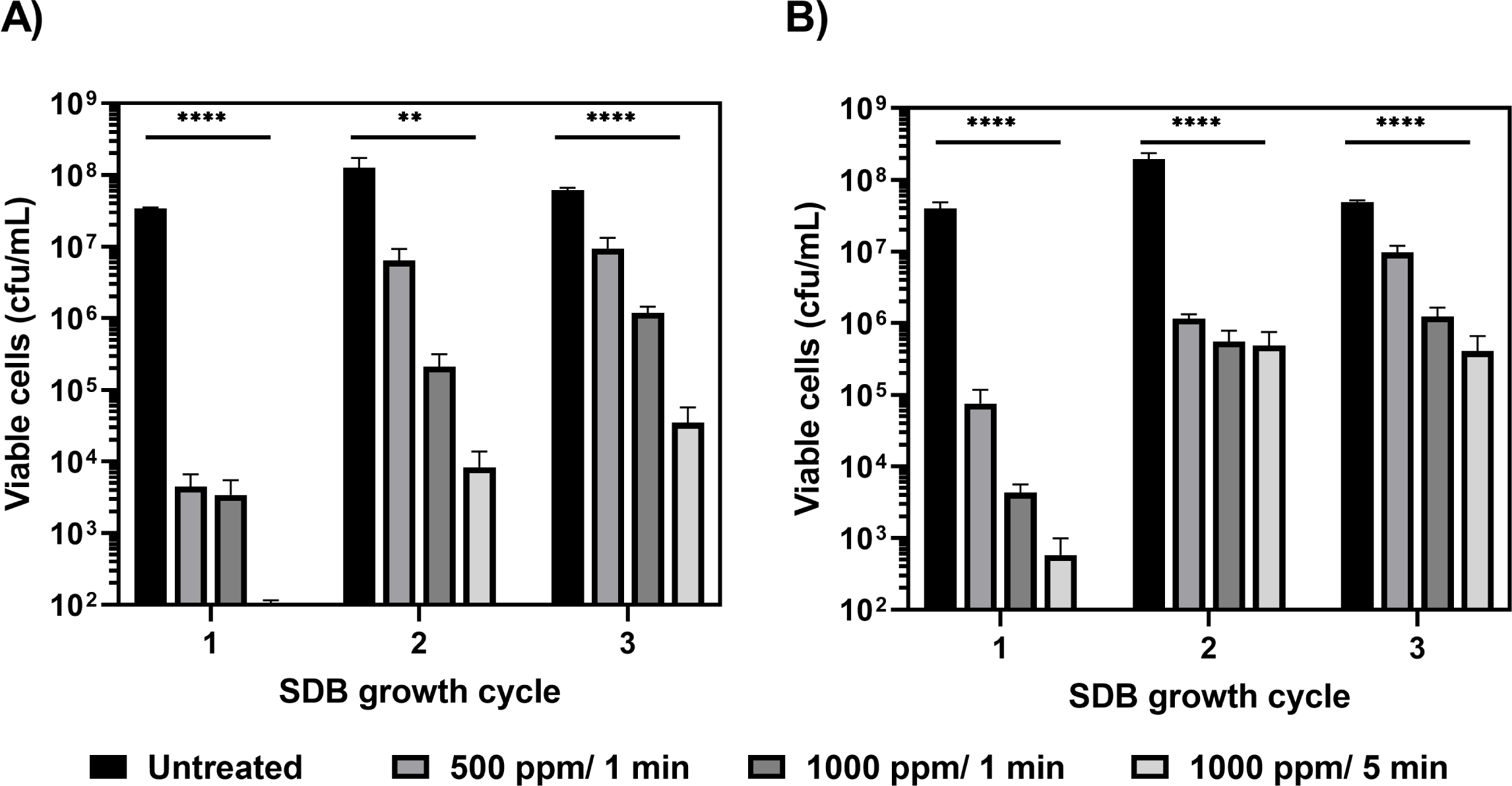
Repeated cycles of wet and dry conditions increase protection against sodium hypochlorite disinfection. *Candida auris* isolates NCPF8973 (A) and NCPF8978 (B) were grown as dry surface biofilms (DSBs) over one, two or three cycles of the protocol. At the conclusion of each cycle biofilms were treated with 500 – 1000 ppm sodium hypochlorite disinfectant for 1 or 5 min or sterile water (untreated) for 5 min. Data represents the mean (± SEM) viable counts for experiments performed in triplicate on three separate occasions. Treatments were compared to untreated biofilms using one-way ANOVA with Benjamini, Krieger and Yekutieli two-stage step-up method for controlling false discovery rate post-hoc analysis: ** p< 0.01, **** p< 0.0001.

### Transcriptional profiling of *C. auris* dry surface biofilms

RNA-sequencing and transcriptomics analysis were performed on cells following planktonic and DSB growth to investigate mechanisms that could contribute to tolerance to disinfection. Total raw reads were >10 million for each sample, and average read alignment was >90%, considered acceptable for transcriptomics analysis (**Table S1**). Principal component analysis (PCA) of expression data demonstrated that the greatest source of variance between samples (PC1) was the growth mode, namely DSB or planktonic conditions, followed closely by the isolate (PC2) as expected (**Fig. 5A**). Differential expression (DE) analysis identified a total of 178 upregulated genes in DSBs relative to planktonic cells, and 169 downregulated genes (**Fig. 5B**). Given that expression appeared to be isolate-specific in the PCA, the number of DE genes between DSBs and planktonic cells was also examined for each individual isolate. In total 227 and 201 genes were found to be up- or down-regulated, respectively, in the single-celled isolate NPF8973 (**Fig. 5C,D**), fewer than the 247 and 239 up- or down-regulated genes, respectively, in the aggregating isolate NCPF8978 (**Fig. 5C,D**).

**Fig. 5.**
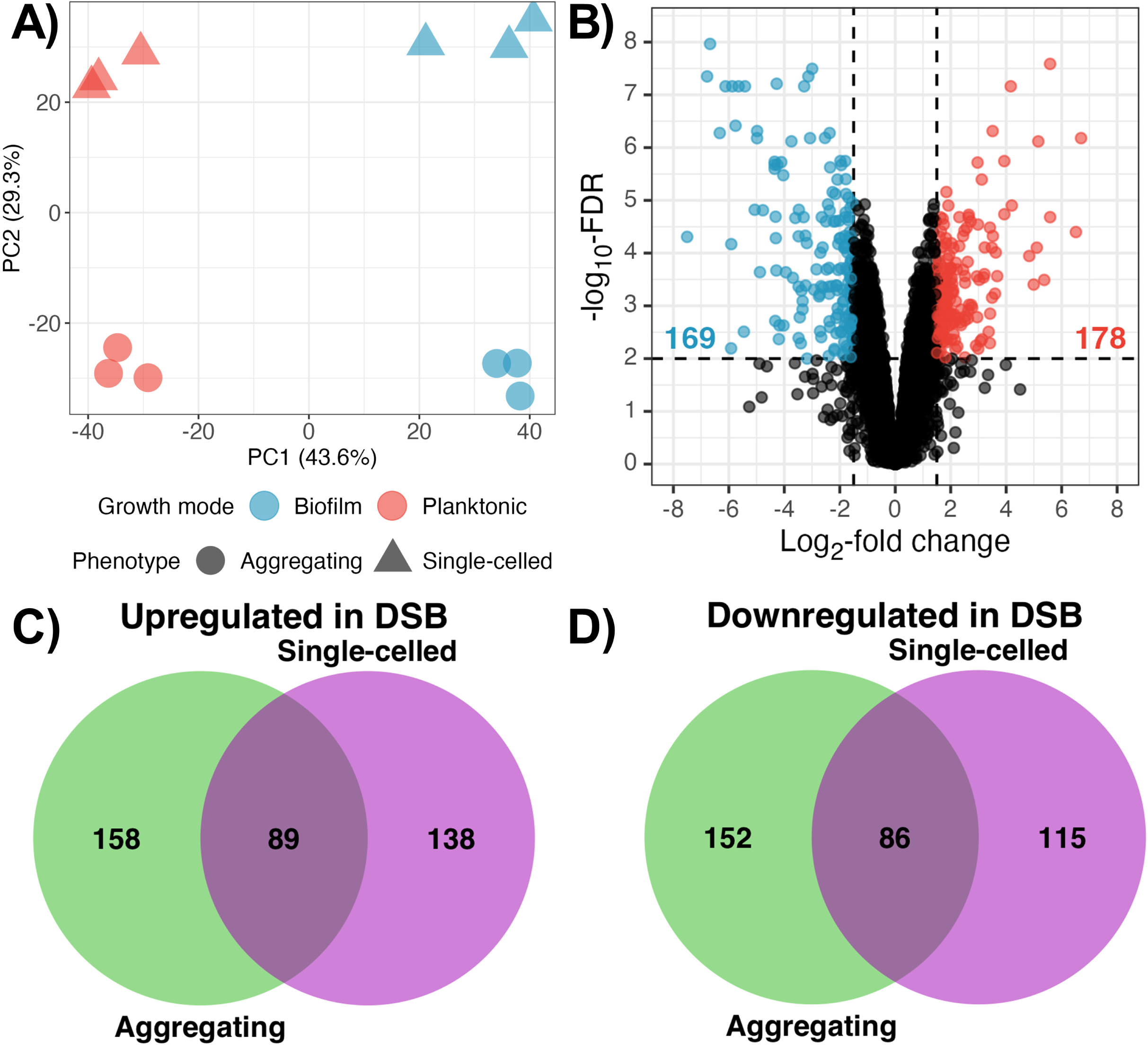
Gene expression by dry surface biofilms and planktonic cells is strain- and growth-mode specific. RNA-sequencing and transcriptional profiling was performed on cells grown for three cycles of the DSB protocol, and compared to planktonic cells. (A) Sources of variability between samples were examined by principal component analysis using filtered and normalised expression data. Principal components 1 and 2, which accounted for over 70% of the variance, were plotted against each other, and samples coded by colour for growth mode, and shape for phenotype. (B - D) Differential gene expression analysis was carried out to determine up- or down-regulation of genes in DSB relative to planktonic cells, based on a log2-fold change in expression of ≥ 1.5 with adjusted p-values <0.01. (B) Volcano plot of fold-changes in gene expression between DSB and planktonic cells, versus probability of differential expression using combined data from both isolates; highlighted boxes show genes that are upregulated (blue) or downregulated (orange) in DSB cells relative to planktonic cells. (C and D) Venn diagrams of upregulated (C) and downregulated (D) genes between DSB and planktonic cells for single-celled isolate NCPF8973 (purple) and aggregating isolate NCPF8978 (green).

Gene ontology (GO) analysis identified 11 enriched terms amongst genes upregulated in DSBs for the combined-isolate analysis (**Fig. 6A**). Nine terms were concerned with ribosomal structure and assembly or translation, and the final two with transmembrane transport. Two ATP-binding cassette (ABC) transporters, *CDR1* and *CDR4*, as well as thirteen as-yet uncharacterised open-reading frames (ORFs), were identified amongst upregulated genes (**Fig. 6B**), supporting the role of these genes in drug and, potentially, disinfectant tolerance. Cytoplasmic iron uptake was also upregulated by DSBs, including the siderophore transporter *SIT1*, ferric reductase *FRE9* and associated iron permease *FTH1*, ferroxidase *FET3/FET31*, and the transcriptional activator *SEF1* (**Fig. 6B**). Seven uncharacterised ORFs thought to be members of the *SIT1* family which is expanded in *C. auris* relative to other *Candida* spp. (23), were also upregulated in DSBs (**Table S2**). This includes the two most highly upregulated ORFs *B9J08_001548* (logFC=8.2) and *B9J08_001547* (logFC=6.7), which remained in the top three-most highly upregulated genes for each isolate, with similar log-FCs (**Table S2**). Such strong and consistent upregulation of putative siderophore transporters highlights iron as a crucial factor in DSB formation by *C. auris*. Other ORFs which are putative orthologues to ABC transporter *SNQ2*, and major facilitator superfamily (MFS) transporters *BSC6* and *HGT2,* were also upregulated (**Table S2**). Genes for the synthesis of ergosterol (*ERG3* and *ERG6*; **Fig. 6B**), and mannans (*MNN26*; **Table S2**), crucial for maintenance of the cell membrane and cell wall, respectively, were also amongst those upregulated by DSBs.

**Fig. 6.**
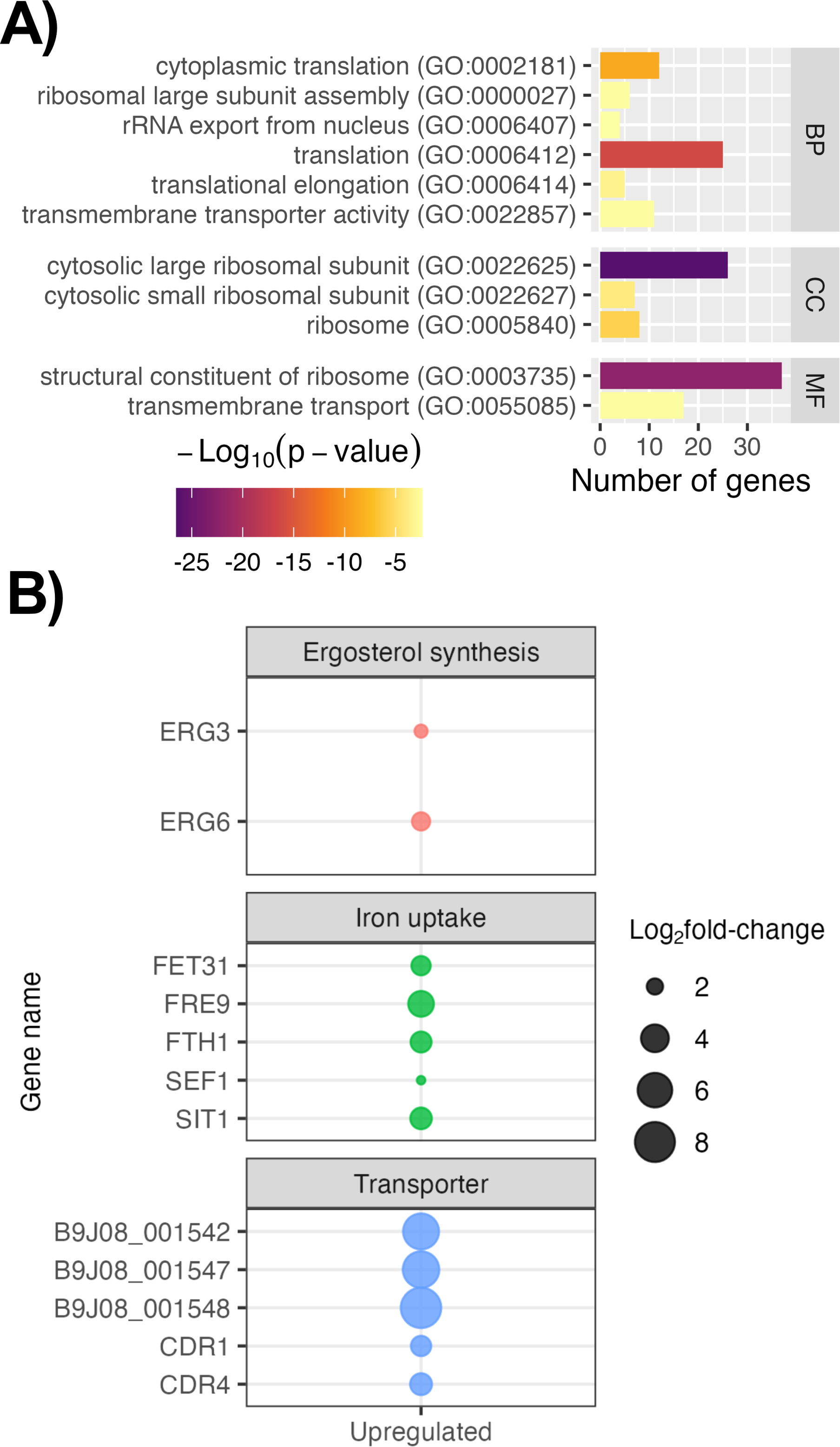
Dry surface biofilms upregulate expression of genes for transmembrane transport and iron uptake. (A) Gene ontology (GO) overrepresentation analysis was conducted on the 179 upregulated genes in DSBs and significantly enriched GO terms (p<0.05, Benjamini-Hochberg FDR multiple comparisons) were returned. (B) Genes of interest within the upregulated subset were further examined and assigned general categories based on their function. BP-biological process; CC – cellular component; MF – molecular function.

Overrepresentation analysis on genes downregulated by DSBs in both isolates found 17 enriched GO terms (**Fig. 7A**), two of which were related to cell wall and its organisation, and the remainder concerned with nuclear and/or cytoplasmic aspects of cell division. In contrast to iron scavenging pathways, a set of genes involved with mitochondrial iron homeostasis, haem biosynthesis, and iron ion binding, *FSF1*, *HEM13*, *HXM1*, *ISA1*, *JLP1* and *NFU1* were found to be downregulated in DSBs (**Fig. 7B**). Cell wall integrity proteins *PGA6*, *PGA26*, *PGA38*, *PGA54*, *PGA58*, *PIR1*, and *DSE1* were all downregulated (**Fig. 7B**). Cell wall remodelling enzymes *FGR41*, *SUN41* and *SCW11* for β-glucans, *CHS1*, *CHT3*, *CRH11* and *GFA1* for chitin, and *ECM331* for mannan deposition, were also downregulated (**Fig. 7B**). *ACE2*, a transcription factor which activates *CHT3* and *SCW11* expression for cell separation, was also downregulated (**Table S2**). Downregulation of genes responsible for nuclear division and associated cell wall remodelling and separation, Not also suggests that the DSBs have reached peak cellular density, consistent with the plateau in viable counts within DSBs following the first cycle of growth (**Fig. 3A**). Collectively, the RNA-seq data presented confirmed that DSBs contain metabolically active cells following the final cell cycle, although cellular division is less likely. Transmembrane transporter activity and iron homeostasis are likely key processes in the formation and/or maintenance of DSBs in *C. auris*, and may play a role in the development of tolerance to NaOCl.

**Fig. 7.**
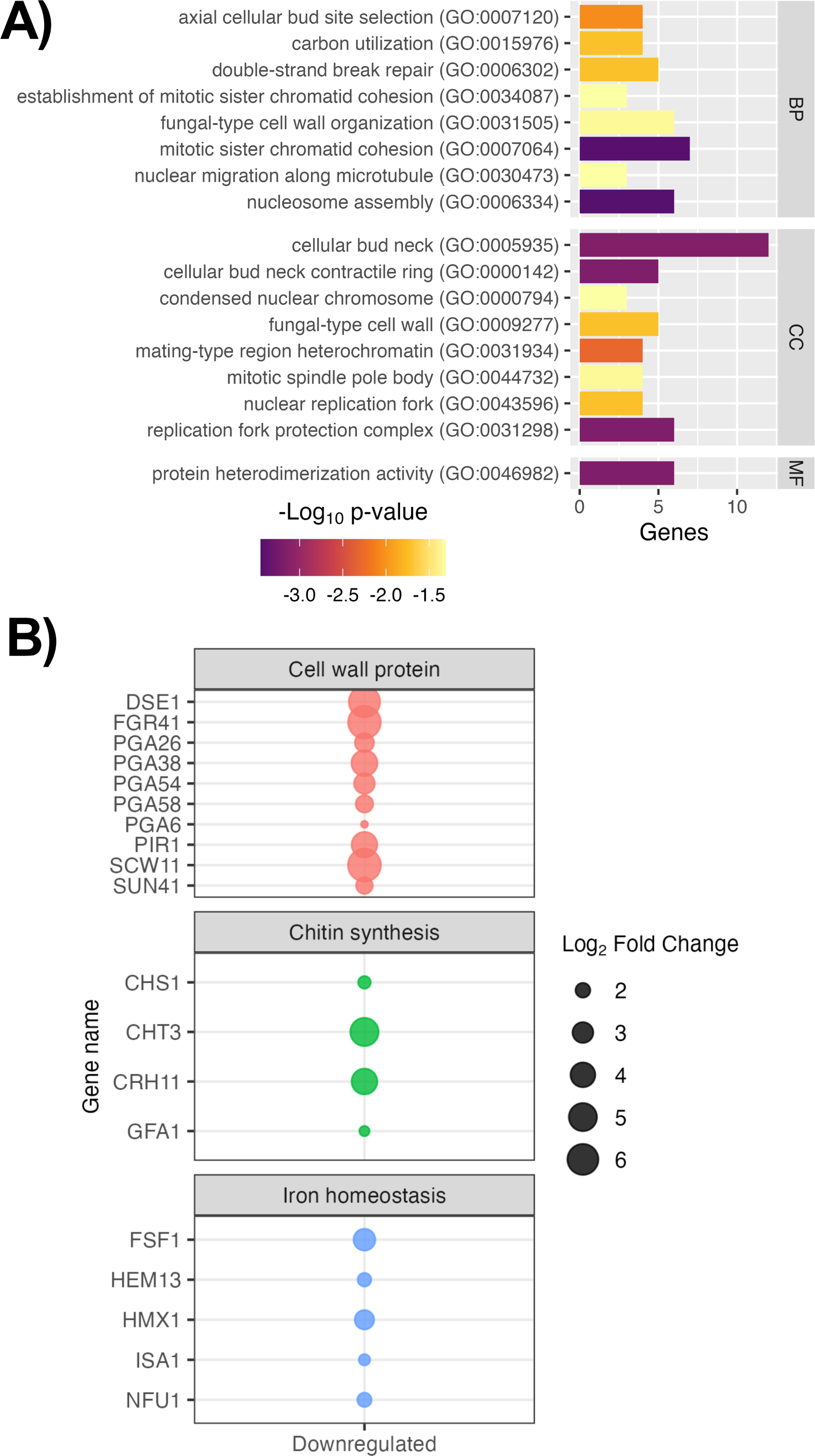
Expression of cell wall remodelling and haem utilisation genes are downregulated in dry surface biofilms. (A) Gene ontology (GO) overrepresentation analysis was conducted on downregulated genes within DSBs and significantly enriched GO terms returned (p<0.05, Benjamini-Hochberg FDR for multiple comparisons). (B) Genes of interest within the downregulated subset of genes were further examined and assigned categories based on their specific function. BP-biological process; CC – cellular component; MF – molecular function.

## DISCUSSION

The global emergence of *C. auris* since 2009 is unlike any other fungal disease (4), and poses a unique challenge within healthcare settings worldwide, where it continues to affect to many vulnerable and at-risk populations (7). Many facilities where *C. auris* is endemic implement proactive screening approaches to detect colonisation and clinical disease, and prevent transmission (24). Whilst such prioritisation has resulted in improved methods for detection of *C. auris*, many protocols for decontamination of colonised skin and abiotic surfaces are largely based on data from bacterial pathogens (21). Failure to optimise disinfection protocols for *C. auris* could prolong outbreaks and exacerbate poor patient outcomes. As such, there is an urgent need for representative testing of disinfectants to guide development and implementation of cleaning and disinfection protocols within healthcare facilities.

In this study we have demonstrated that clinical isolates of *C. auris* are intrinsically sensitive to killing by NaOCl, using two models of suspension testing. Disinfection efficacy studies against planktonic cells typically demonstrate that *C. auris* isolates are sensitive to the chemical activity of chlorine-based agents at concentrations ranging between 100-6500 ppm, depending on contact time (13–15, 18–20). Contact times between 1-5 min achieve total eradication of viable cells using 100-1000 ppm chlorine-based agents (14, 15, 18), consistent with our findings of ≥6-log_10_ reductions with 500-1000 ppm NaOCl for the same time. When cultures are dried on surfaces, however, minimal log_10_-reductions are observed with 500-2000 ppm NaOCl for 1 min (19), and concentrations of NaOCl ≥3900 ppm are required to achieve a minimum 4-log_10_ reduction in cells with 1 min contact time (13, 20, 25). It is thus likely that drying of biological material and/or surface attachment such as biofilm formation, facilitates tolerance to disinfection in *C. auris* cells as is shown with other infectious agents.

The seminal study by Ledwoch & Maillard (22) previously characterising DSB formation by *C. auris* reported development of sparse, monolayered biofilms lacking significant ECM. These biofilms were effectively removed by mechanical disinfection with 1000 ppm NaOCl wipes, representing ≥7-log_10_-reduction in fungal burden. The isolate used in their study, however, has since been shown be more susceptible to NaOCl disinfection than other *C. auris* isolates (26), and clade II isolates are neither associated with outbreaks nor antifungal resistance (27). In contrast, isolates from outbreak-associated clades I and III used in our study formed thick, multilayer DSBs with extensive ECM over three cycles of growth. DSBs development coincided with tolerance to the chemical activity of NaOCl, which was demonstrated by increased survival following successive cycles of growth. This tolerance occurred despite plateaued numbers of cells within DSBs after each cycle, further corroborated by downregulation of genes associated with nuclear replication and cell separation such as *ACE2* (B9J08_000468), *SCW11* (B9J08_003120) and *CHT3* (B9J08_002761). *ACE2* is a transcriptional activator of a number of genes including *SCW11*, *CHT3* and *DSE1* which were all downregulated in DSBs. Interestingly, disruption of *ACE2* has been previously shown to induce cell separation defects resulting in aggregation (28), in agreement with our data this potentially suggests DSB formation occurs concurrent with aggregation induced by environmental stress and starvation (29).

The role of efflux pumps in establishing tolerance and/or resistance to antimicrobials has been widely documented in both bacterial and fungal species (30). Multiple transporters belonging to ABC (*Cdr1* and *Snq2*) and MFS (*Mdr1* and *Flu1*) protein families are expressed by different *Candida* spp., and accept a wide variety of structurally diverse xenobiotics as substrates. *In vitro* models of *C. auris* have demonstrated that constitutive expression of *CDR1* and *MDR1* by both planktonic cells and biofilms is strongly associated with increased MICs to azole antifungals (6, 31). Furthermore, deletion of *CDR1* but not *MDR1*, restored clinical sensitivity to fluconazole and itraconazole in resistant *C. auris* isolates, highlighting *CDR1* as a key player in tolerance to small-molecule antifungals (31). We also observed upregulation of *CDR1* (B9J08_000164) and *CDR4* (B9J08_000479), as well as ORFs with predicted transporter activity, by DSBs in our study, although a role for efflux pumps in NaOCl tolerance is unclear. Clinically relevant bacteria such as *Mycobacterium* spp. possess a redox-sensing mechanism whereby oxidation of transcriptional repressors by intracellular NaOCl results in expression of efflux pumps, leading to subsequent removal of NaOCl molecules from cells (32). Given that similar mechanisms have also been found in *Pseudomonas*, *Legionella* and *Bacillus* spp.(33), it is likely that *CDR1* and *CDR4* efflux pumps which are already expressed within *C. auris* DSBs facilitate removal of NaOCl and enable tolerance. Antifungal tolerance in *C. auris* biofilms also occurs due to the ECM components such as mannans, which Dominguez and colleagues (5) previously demonstrated actively sequesters fluconazole molecules away from cells, thereby preventing its activity against cells. In our model, *MNN26* (B9J08_004650) for mannan synthesis was upregulated, whilst *ECM331* (B9J08_004382) required for cell wall mannan deposition was downregulated, potentially suggesting that mannan synthesis was utilised extracellularly.

Iron homeostasis is critical to maintain intracellular stores of the ions, which act as cofactors for many proteins and enzymes within cells (34). Low iron conditions induce expression of *SEF1* and *HAP43* transcriptional factors in *C. albicans* and *C. parapsilosis*, which in turn activate genes related to iron uptake and scavenging including *FET31*, *FTH1, FRE9*, and *SIT1*, and turns off iron usage genes such as *HMX1*, and *HEM13* to main intracellular iron levels (34, 35). Biofilm formation and virulence are also closely linked with iron homeostasis, as *SEF1* is required to maintain adhesion in *C. albicans* (36). In our study, similar expression patterns were observed for the aforementioned genes, suggesting conserved function of iron in maintaining *C. auris* biofilms.

Taken together, our data demonstrate that *C. auris* isolates from outbreak-associated clades are capable of developing robust biofilms under the DSB protocol. Extended growth under these conditions results in enhanced tolerance to NaOCl disinfection, resulting in loss of efficacy. Upregulation of small-molecule efflux pumps and interference of the biofilm matrix are likely mechanisms by which *C. auris* DSB cells gain tolerance to NaOCl disinfection. Further functional studies into transmembrane efflux pumps such as *CDR1*, and iron homeostasis in DSBs are needed to elucidate mechanisms behind biofilm-associated NaOCl tolerance which facilitate the persistence of this robust organism within the healthcare environment. Our work highlights the importance of selecting appropriate, clinically representative models to test efficacy of disinfectants against nosocomial pathogens.

## MATERIALS AND METHODS

### Microbial culture and standardisation

Clinical isolates of *C. auris* from two outbreak-associated phylogenetic clades, that were not multi-drug resistant, displayed varying biofilm forming capacity, and included aggregating versus single-celled phenotypes, were selected for this study (**Table 1**) (37). Isolates were maintained on Sabouraud’s dextrose agar (SAB [Sigma-Aldrich]) following incubation for 48 h at 30°C. Overnight broths were prepared by inoculation of colonies into yeast peptone dextrose medium (YPD [Sigma-Aldrich]) and incubation at 30°C for 18 h in an orbital shaking incubator at 150 rpm. Cells were quantified using a Neubauer haemocytometer, and standardised to the required concentration in appropriate medium as detailed in subsequent sections.

**Table 1.** Key characteristics of selected *Candida auris* strains.

| Growth phenotype | Isolate | Biofilm former phenotype | Phylogeographic clade |
| --- | --- | --- | --- |
| **Single-celled** | NCPF8990 | High | I |
|  | NCPF8973 | Moderate | I |
|  | Strain 174 | Low | I |
| **Aggregating** | NCPF8991 | High | I |
|  | NCPF8978 | Moderate | III |
|  | Strain 182 | Low | III |

### Dry surface biofilm assay

A dry surface biofilm (DSB) protocol was adapted from Ledwoch & Maillard (22). The DSB protocol consisted of three cycles of alternating growth under hydrated conditions for 48 h, followed by a further 48 h under dry conditions (**Fig. 1**); all incubation periods were carried out at room temperature. Isolates were standardised to 1×10^6^ cells/mL in Roswell Park Memorial Institute (RPMI) media supplemented with 5% (v/v) foetal calf serum (FCS [Sigma-Aldrich]) to simulate organic load. Growth under hydrated conditions was initiated by seeding cells in cell-culture treated 24-well polystyrene microtitre plates (Thermo Fisher Scientific). The subsequent dry phase consisted of removal of all liquid from wells. Biofilms were washed once with sterile phosphate-buffered saline (PBS [Oxoid]) to remove non-adherent cells after the first hydration cycle. Characterisation of biofilms, including quantitation of viable cells and total biomass, and disinfection efficacy testing, was carried out after each dry phase, as described below. Where appropriate, biomass was harvested from plates by addition of PBS and scraping the surface using a sterile pipette tip.

### Disinfectant preparation

NaOCl disinfectant was purchased as a solution containing 5% or 50,000 ppm active chlorine (Acros Organics) and stored at 4°C until use. Working solutions of NaOCl were prepared fresh for each experiment by dilution in RPMI Sigma-Aldrich or sterile water to concentrations between 4-4000 ppm.

### Disinfectant sensitivity testing

Planktonic minimum inhibitory concentrations (PMICs) were determined for NaOCl adapted from the CLSI-M27 broth dilution method (38). In brief, NaOCl was serially diluted two-fold in wells of 96-well round bottom microtitre plates between 7.8 and 4000 ppm in RPMI, at double the final concentration. Planktonic cells were added to wells at a final concentration of 1×10^4^ cells/mL in RPMI, and plates incubated at 37°C for 24 hr. Microbial growth was assessed visually, and by measuring the optical density at 530nm using a spectrophotometric plate reader. The MIC was determined as the concentration of NaOCl required to inhibit 50% or 90% of microbial growth relative to untreated controls.

### Disinfection efficacy protocol

The efficacy of NaOCl disinfection against planktonic and biofilm cells was determined according to the method by Short *et al.* (2). In brief, planktonic cells were initially standardised to 2×10^8^ cells/mL in PBS + 5% FCS and aliquoted into 24-well microtitre plates for testing. Biofilms were also grown in 24-well microtitre plates for one, two, or three cycles of the DSB protocol. NaOCl was diluted from stock to 500 to 2000 ppm in sterile water, and cells and biofilms treated with a final concentration of 500 ppm or 1000 ppm for 1- or 5-min. Sterile water was used as a positive control (untreated). The active agent was neutralised with sterile sodium thiosulfate solution (Na_2_S_2_O_3_; 5% w/v in H_2_O [Sigma-Aldrich]) added to all samples for 15 min.

Planktonic cells were treated with an equal volume of 2x NaOCl to cells and neutralisation carried out using two-volumes Na_2_S_2_O_3_. In contrast, 1x NaOCl was added to directly to washed biofilms, and neutralised with an equal volume of Na_2_S_2_O_3_. Surviving cells were resuspended in fresh PBS following neutralisation, for quantification as detailed below.

### Viable cell counts

The Miles and Misra technique (2) was used to quantify viable cells during formation of DSBs or following disinfection of planktonic cells and DSBs. Cells and biomass were collected and serially diluted 10-fold in sterile PBS, and cultured onto SAB plates in triplicate across each dilution. Plates were incubated for 48 h at 30°C. Determination of colony forming units (CFU) per mL was performed using the average colony count of replicates.

### Quantitation of total biofilm biomass

Total biomass was measured at the end of each growth cycle using the crystal violet assay (39). Washed DSBs were stained with 0.5% (w/v) crystal violet for 15 min, after which excess was removed and biofilms washed twice. Plates were dried before eluting bound dye with absolute ethanol, of which 75 μL was transferred to a 96-well microtitre plate and the optical density measured at 570 nm in a plate reader.

### Scanning electron microscopy

DSB ultrastructure was imaged using scanning electron microscopy (SEM) (40). Briefly, biofilms grown for one or three cycles on Nunc^TM^ Thermanox^TM^ coverslips (Fisher Scientific), were fixed using 2% paraformaldehyde, 2% glutaraldehyde, 0.15% alician blue power and 0.15 M sodium cacodylate. Samples were counterstained using uranyl acetate with subsequent gradient dehydration in ethanol (30 to 100%). Dried samples were mounted, sputter-coated using gold/palladium and visualised using an IT 100 SEM machine at 1000x and 10,000x magnification (JEOL Ltd).

### RNA extraction and sequencing

A total of 5×10^8^ planktonic cells from isolates NCPF8973 and NCPF8978 were collected by centrifugation. DSBs were also grown for each isolate as detailed above and biomass harvested by scraping and centrifugation. RNA was extracted from samples using the RiboPure Yeast kit (Invitrogen) according to manufacturer’s instructions. The yield and quality of RNA was assessed using a DS-11 Fx+ spectrophotometer (DeNovix, USA), and the RNA Integrity Number (RIN) determined by Bioanalyser (Agilent). All samples had a yield >500 ng/μL and RIN ≥7.2, deemed acceptable for downstream applications. RNA sequencing (RNA-seq) was performed by Novogene (https://www.novogene.com/) on a Novaseq 6000 platform (Illumina) to produce 150 bp paired-end reads according to their standard protocols. The data presented in this publication have been deposited into NCBI’s Gene Expression Omnibus (https://www.ncbi.nlm.nih.gov/geo/) and are accessible through GEO Series accession number GSE239851.

### Transcriptomics analysis

Transcriptional profiling using RNA-seq data was carried out using R (v4.3.0) in RStudio (v32), and all packages used within are open-source and available through the Comprehensive R Archive Network (https://cran.r-project.org), Bioconductor (https://bioconductor.org), or GitHub (https://github.com). Data handling and graphical outputs were performed using tidyverse packages (v2.0.0). Genomic sequences, gene annotations, and gene ontology (GO) annotations for *C. auris* reference strain B8441 were accessed from the Candida Genome Database (v s01-m01-r31; http://www.candidagenome.org) and the GO Consortium (http://geneontology.org). Raw sequencing reads were quality-controlled using FastQC (v0.12.1). Genome indexing and read-mapping were carried out in Kallisto (v0.48.0). Abundance outputs were converted to log_2_-transformed counts per million (logCPM), filtered, and normalised to the library size using EdgeR (v3.42.4).

Differentially-expressed genes were selected based on log_2_-fold change ≥1.5 (logFC) in either direction and a false discovery rate (FDR)-adjusted significance p <0.01, determined using limma-voom (limma v3.56.1). GO overrepresentation analysis (hypergeometric distribution, FDR multiple comparisons p <0.05) was performed on gene sets containing up- and down-regulated genes for each analysis using clusterprofiler (v4.8.1).

### Data analysis

Data were analysed in Microsoft Excel (v16.73) and graphs compiled in GraphPad PRISM (v9.1.0). Normality tests were performed prior to appropriate statistical analysis. Viable cells from disinfected biofilms were compared to untreated biofilms using one-way ANOVA with Benjamini, Krieger and Yekutieli two-stage step-up method for controlling FDR as post-hoc analysis. Significance was set at p <0.05.

## ACKNOWLEDGEMENTS

We would like to acknowledge the funding support of Haleon and the BBSRC Industrial CASE PhD studentship for M. B. (BB/V509541/1). We would also like to thank Margaret Mullin at the Glasgow Imaging Facility for her expertise with scanning electron microscopy.

---

**Fig. S1. *Candida auris* clinical isolates show heterogeneity in biofilm forming ability.** Twenty-six clinical isolates were screened for biofilm formation. Cells standardised to 1×10^6^ cells/mL in RPMI were seeded into wells of 96-well microtitre plates and biofilms grown over 24-hr. Total biomass from washed biofilms was detected by staining with 0.5% (w/v) crystal violet solution and bound dye eluted in ethanol before quantification at 570 nm. Dotted and dashed lines represent cut-off absorbance values for classification as high or moderate biofilm formers, respectively. Blue and green samples represent isolates which were selected for use in this study.

